# Large-scale recovery in Costa Rica’s payment for ecosystem service program

**DOI:** 10.1101/2024.09.03.610944

**Authors:** Giacomo L. Delgado, Johan van den Hoogen, Daisy H. Dent, Tom Bradfer-Lawrence, Leland K. Werden, Rebecca Cole, Cristian Diaz Quesada, Jose-Angel Jimenez Fajarado, Alberto Méndez Rodríguez, Eduardo Mesén Solorzano, Gilmar Navarrete Chacón, Mario Coto, Irene Suarez Perez, Lucas Vahlas, Yuting Liang, Thomas Ward Crowther

## Abstract

Costa Rica implemented the world’s first national-scale Payment for Ecosystem Service (PES) program in 1996 and now protects over 200,000 hectares. By distributing wealth towards local land-stewards, Costa Rica’s program has helped to limit deforestation at a national scale, but the large-scale ecological implications have yet remained unclear. Here, we use a massive ecoacoustic dataset to evaluate how this program has impacted the ecological integrity of PES forests across the entire Nicoya Peninsula. At the times and frequencies that are indicative of native biological activity, we reveal dramatic increases in the integrity of PES soundscapes, relative to those of natural protected areas. Specifically, natural regeneration sites were 97.79% more acoustically similar to reference forests (absolute mean similarity of 0.539) than they were to disturbed pastures, while acoustic recovery of plantations lags behind (79.66%; 0.489). These findings are strongly suggestive of large-scale ecological recovery, constituting some of the most robust evidence to date that restoration initiatives can benefit biodiversity on large spatial scales.

**Study overview:** Costa Rica’s PES program pays landowners to encourage forest recovery and compensate them for opportunity costs. Most payments subsidize land ‘conservation’, in which participants allow existing forests to naturally regenerate. Some payments are also offered to produce timber through ‘plantations’, which are often monocultures of exotic tree species. Despite the program’s importance to Costa Rica’s conservation efforts, little is known about whether these forest systems are recovering their natural characteristics. To investigate the dynamics of the PES program, we recorded continuous 6-day soundscapes in 119 sites across the Nicoya Peninsula of Costa Rica (Supplementary Figure 1). Specifically, we characterized the soundscapes across 4 land-use types: (i) 19 reference pastures, (ii) 43 PES monoculture tree plantations, (iii) 39 PES natural regeneration sites, and (iv) 18 reference forests. Sites from each land-use type are distributed across the Nicoya Peninsula’s climate and edaphic gradient, allowing us to capture substantial variation in ecological outcomes. We determined the areas of acoustic space where most animals vocalize and where ecological responses to recovery were most likely to be detected (Figure 1). We then identified how and to what extent the soundscapes of natural regeneration and plantation sites had changed over the last 27 years. We find evidence that naturally regenerating forests within the PES have recovered substantially when compared to reference forests, while plantation systems lag behind (Figure 3). Our findings reaffirm the importance of ecosystem conservation, while suggesting redistributive policy mechanisms can accelerate nature protection at scale.

## Introduction

Land degradation is the largest global driver of biodiversity loss, causing more than 10% of global emissions and directly threatening the livelihoods of at least 3.2 billion people^1^. Implementing mechanisms to protect ecosystems and prevent the loss of biodiversity at scale have proved among this century’s most complex social and ecological challenges^2^. A growing body of evidence has tied the degradation of Earth’s ecosystems to the inequitable distribution of wealth^3,4^, which leads to overconsumption by the hyper-rich^5^ and forces the poor into the most degraded margins of the economy and environment^4,6^. As such, recent studies have proposed that mechanisms that combat this inequality, by driving equitable wealth redistribution towards local land stewards have the potential to reverse this trend and enable large-scale ecological recovery^7,8^. Despite strong conceptual support for the link between equitable wealth distribution and environmental recovery, testing this hypothesis remains challenging because we lack empirical evidence that pairs long-term redistribution with tangible environmental outcomes.

Over recent decades, various policy mechanisms have been implemented to finance the conservation and restoration of nature, with mixed results. One promising mechanism to promote the distribution of wealth for the protection of nature are Payment for Ecosystem Service (PES) programs, for which Costa Rica is recognized as a global leader. The country established the world’s first national-scale PES program in 1996^9^ and now protects over 200,000 hectares with the specific goals of recognizing the water provisioning, carbon storage, biodiversity protection and scenic ‘beauty’ of the nation’s forests^10^. While the financial scale of program subsidies^11,12^ is limited, (∼70USD/ha/year) and research has found mixed effects on poverty alleviation^13,14^, the program’s implementation represents a unique national-scale experiment to evaluate the environmental impacts of distributing wealth directly to local land stewards at scale.

Alongside a wide suite of regulations and socio-economic conditions^15,16^, the PES program has made Costa Rica a model in environmental protection. Previous studies employing satellite observations suggest that the PES has directly helped to increase forest cover^17^, carbon storage and, potentially, habitat for endangered species^18^. Yet, despite the need to track progress on biodiversity targets^19^ collecting standardized monitoring data at large spatial scales remains a key challenge. In the absence of standardized ground-sourced data collection, the implications for ecological integrity within the PES remain untested.

Traditional biodiversity surveys are often time consuming and expensive, which limits their application to specific geographies with limited their scalability^20^. To overcome these challenges, recent studies have highlighted the potential of passive-acoustic monitoring techniques as scalable solutions with the potential to track changes in biodiversity and ecological condition^21,22,^. While caution is requited for interpreting information about the richness or diversity of species, well-designed studies based on strong theoretical understandings^23,24^ of “ecoacoustic” indices^25^can provide insights into the integrity of the acoustic ecosystem (i.e., soundscapes), which can serve as indicators of ecological recovery^26^. Yet until now, few studies have utilized large acoustic datasets to detect ecosystem-level ecological changes at this scale and none evaluate the outcomes of multiple paired long-term conservation and restoration interventions.

Here, we use ecoacoustic recordings from 119 sites across a broad geographic extent to test whether the increases in Costa Rica’s forested area have been accompanied by other aspects of ecological recovery. We characterize soundscapes of 119 sites along a spectrum from highly to minimally disturbed. ‘Reference pasture’ sites – where natural forests have been removed – reflect these landscapes before restoration. At the other end, ‘reference forests’ sites represent the state of natural forest ecosystems outside the PES parcels and have been protected in the long-term by entities such as the National Park system (SINAC). These two baseline states are used to contextualize the dynamics of two common program interventions: PES restoration with natural regeneration (∼30-60 years old) and PES restoration with timber plantations (∼7-20 years old). This approach enabled us to assess how the soundscapes of forests within the PES have developed as a result of the program. Taken alongside existing data on forest cover and carbon storage, these results provide the first assessment of the success of this redistributive program in catalyzing national-level ecological recovery.

### Tailoring the Acoustic Space

A key challenge in biodiversity monitoring stems from the difficulty in assessing the dynamics of ecological systems in a standardized way. The acoustic signature, or soundscape, of an ecosystem is rich in both ecological^25,27^ and functional^28,29^ information but the complexity of sound presents significant interpretive challenges. Acoustic indices were developed as metrics to simplify this complexity^30^. Many indices were developed as proxies for biological diversity, but associations are often weak or contradictory^22,31^, limiting their applicability. Advances in recording technology make collecting massive datasets increasingly feasible but, in turn, require rapid analysis making the use of acoustic indices more attractive despite limitations. While an increasing number of soundscape ecologists acknowledge that applying indices contextually to relevant areas of the acoustic space may address weaknesses^24,32^, few studies quantify the time and frequency components of soundscapes within their study systems despite promising initial results^33,34^.

Using a data-driven approach we restrict our analysis to the most biologically relevant areas of acoustic space to increase interpretive power (Figure 1). Thus, rather than attempting to capture biological diversity per se across land-use types, we use a modification of the ‘Power Minus Noise’ statistic (PMNsp)^35^ to characterize soundscapes. This statistic can be thought of as describing an aspect of a soundscape’s energy or ‘loudness’. Specifically, ‘power’ is the maximum amplitude in each 93.75Hz frequency band of a spectrogram, after the removal of background ‘noise’ (i.e., the modal amplitude in that band). We modified this metric to give ΣPMN, the sum of all noise-subtracted amplitude values across each 0.01 second cell within each frequency band and minute. This provides an index describing the total intensity of acoustic energy for every 1-minute and 93.75Hz. Higher ΣPMN values therefore indicate loud and dynamic areas of acoustic space, with energy that routinely rises above background noise levels. Low values indicate silence or highly saturated soundscapes where the modal and max amplitude approach each other (e.g., heavy wind or rain).

**Figure 1:**
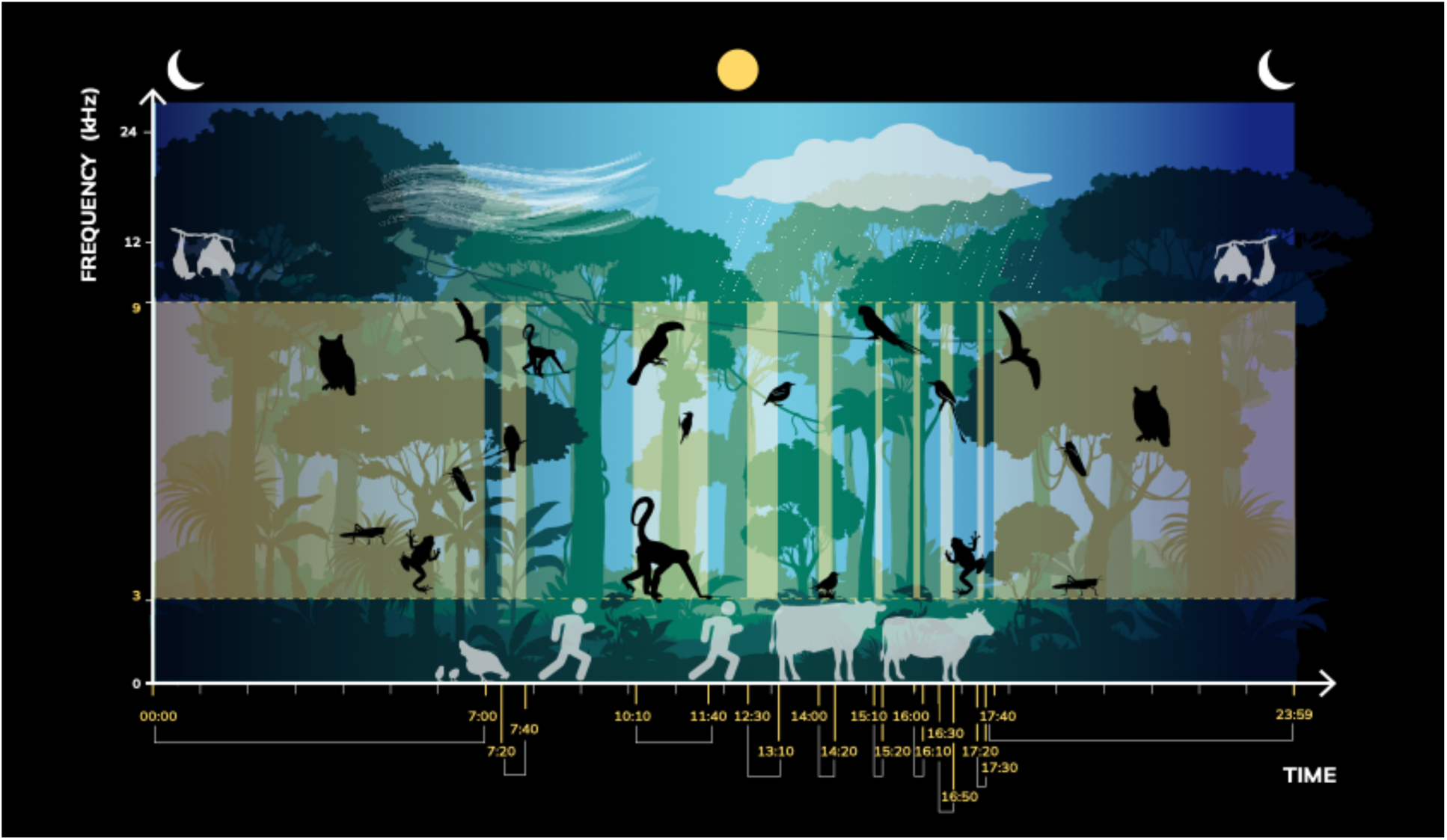
Tailoring the acoustic space to improve acoustic analyses. Soundscape analyses can be improved by identifying the times and frequencies associated with the dynamics of interest. Using manual annotations and modelling approaches we determined that differences in the activity of native fauna, shown here in black, were best understood within frequencies 1 – 9 kHz (on the y-axis) and the times listed along the x-axis in yellow. Outside of this region of acoustic space, differences between land-use types are better explained by climatic differences or other acoustic events (shown in grey).

To understand the temporal dynamics in our dataset we summed ΣPMN across all recorded frequencies (24,000 Hz) and calculated ten-minute means across all sites within land-use types to visualize 24-hour soundscape patterns (Supplementary Figure 3a). Using the Friedman test for repeated measures, we detected significant soundscape differences among all land-use types (x^2^ = 320; p < 0.001). To test the effects of a set of predictive variables on ΣPMN, we employed two modelling approaches: gradient boosted machines and linear-mixed models which were fitted individually to each time-bin, using the Akaike Information Criterion (AIC)^36,37^. Results from both approaches suggest that ΣPMN is strongly predicted by land-use type (Supplementary Figure 2), indicating that our index captures land-use type specific ecological signatures. Time-bins in which the selected linear-mixed models retained land-use type as a predictive variable enabled us to identify the times of day in which these relationships were strongest (Figure 1). Specifically, these include all nocturnal, dusk and dawn hours (i.e., 30 minutes before and after sunrise and sunset respectively) between 17:20 and 7:40 (excluding two 10-minute bins) and various smaller ranges during the day (10:10-11:40, 12:30-13:10, 14:00-14:20, 15:10-15:20, 16:00-16:10 and 16:30-16:50). Differences in soundscape patterns outside of these times are better explained by other site-specific attributes like climate or elevation, therefore we removed these from subsequent analyses comparing land-use types.

To identify the frequency-bands occupied by taxonomic groups of interest (mammals, amphibians, birds, and insects), we manually annotated 120 minutes of recordings across all land-use types (i.e., 30 minutes from each) and the entire diel cycle (i.e., 15 diurnal and 15 nocturnal minutes for each land-use type). This resulted in 8,543 sound types (sonotypes) identified to the taxon level. As predicted by the Acoustic Niche Hypothesis^29^, there was significant frequency partitioning between the four taxonomic groups of interest (ANOVA; p < 0.01): mammals (90% of calls between 318 and 1036 Hz), amphibians (1132 and 4895 Hz), birds (1261 and 8004 Hz) and insects (4831 and 12321 Hz) (Supplementary Figure 3b). As 90% of all biophony (i.e., biological sound) fell within a range of 638 and 8,721 Hz, we consider 0 – 9 kHz to be the biologically active range within our dataset. However, anthropophony (sounds of human origin) and calls from domestic animals made up the majority (68%) of identified low frequency sounds in the 0 – 1 kHz range. The significant overlap with wild mammals in these frequencies, limits the interpretability of results for this low range. As such, all further analyses include only frequencies from 1-9 kHz (Figure 1).

### Soundscape Similarity Tracks Recovery

Within our focal region of acoustic space (i.e., 1 – 9kHz during the time bins identified above; Figure 1), ΣPMN describes biological activity of vocalizing animals such that soundscape similarity to reference forests can function as a proxy for ecological recovery. Well-designed comparisons among soundscapes have been shown to underlie differences in species assemblages and beta diversity^31,36^. Using Wasserstein (i.e., earth mover’s) distances^38^, we calculated acoustic similarity among land-use types for all 880 identified 10-minute and 1 kHz, time-frequency intersections (Figure 2). These values represent both the magnitude and the distance required to move the distribution of mean ΣPMN values (each point representing a single site) of one land-use type, into the distribution of another (Figure 2b). We report these as scaled similarity values such that values of 1 indicate complete overlap (i.e., no distance) and values close to 0 indicate increasingly large acoustic distances. The Friedman statistical test for repeated measures and Wilcox post-hoc tests were used to test for significant differences in similarity among land-use types.

**Figure 2:**
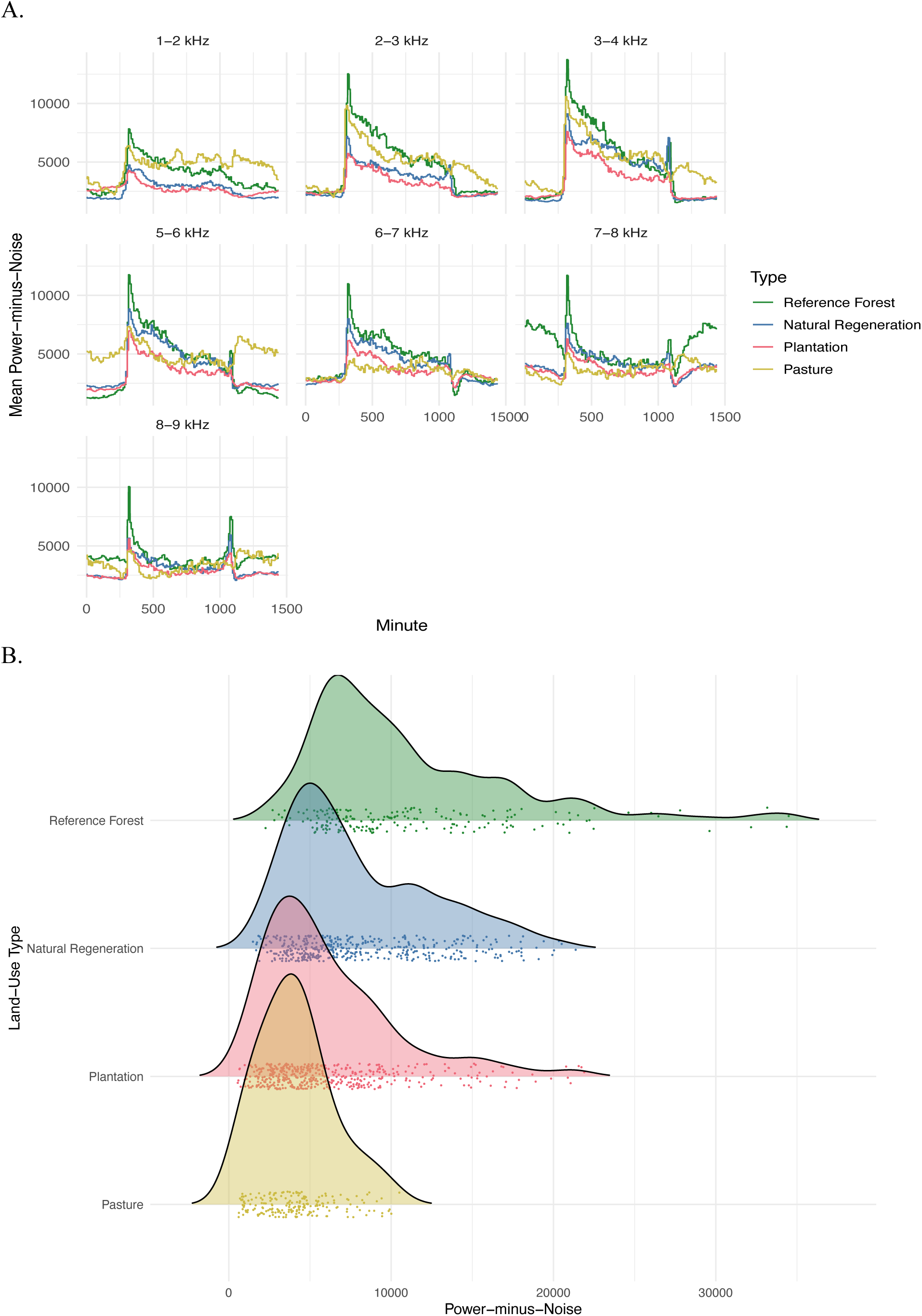
Analyzing acoustic patterns among land-use types requires time-frequency specific comparisons. (A) Patterns in SPMN are highly frequency specific. Mean 24-hour cycles for each land-use type are plotted for the frequency bands included in our analyses. (B) We calculate Wasserstein distances among land-use types within time-frequency intersections. These values represent the distance multiplied by the magnitude required to move one distribution into another. A single set of distributions, colored by land-use type, are shown for the 3 – 4 kHz and 5:15 – 5:25am intersection. Points represent the mean SPMN value for a single minute and site within this intersection.

To understand the status of PES soundscapes, we began by comparing our baseline ecosystems: reference pastures (i.e., most disturbed) and reference forests (i.e., minimally disturbed). As expected, reference forests and pastures had the least similar soundscapes among all land-use types (mean similarity 0.413; Figure 3a). Temporally, differences were most pronounced in the time-bins associated with the dawn (mean similarity of 0.224, minimum of 0.07) and night (mean of 0.415). Alongside low similarity values in the 5-6 and 7-8 kHz bands, these results are consistent with findings that heavily disturbed tropical pastures exhibit lower diversity and altered composition of bird^39,40^ and insect^41,42^ communities. Instead of a strong and sharp peak in acoustic activity at dusk, pastures exhibited a flattened peak that occurred later and lasted longer (Figure 3a), leading to low similarity values with reference forests stretching into the early night. Such loss of acoustic temporal structures has been previously associated with heavily modified ecosystems^33,43^.

**Figure 3:**
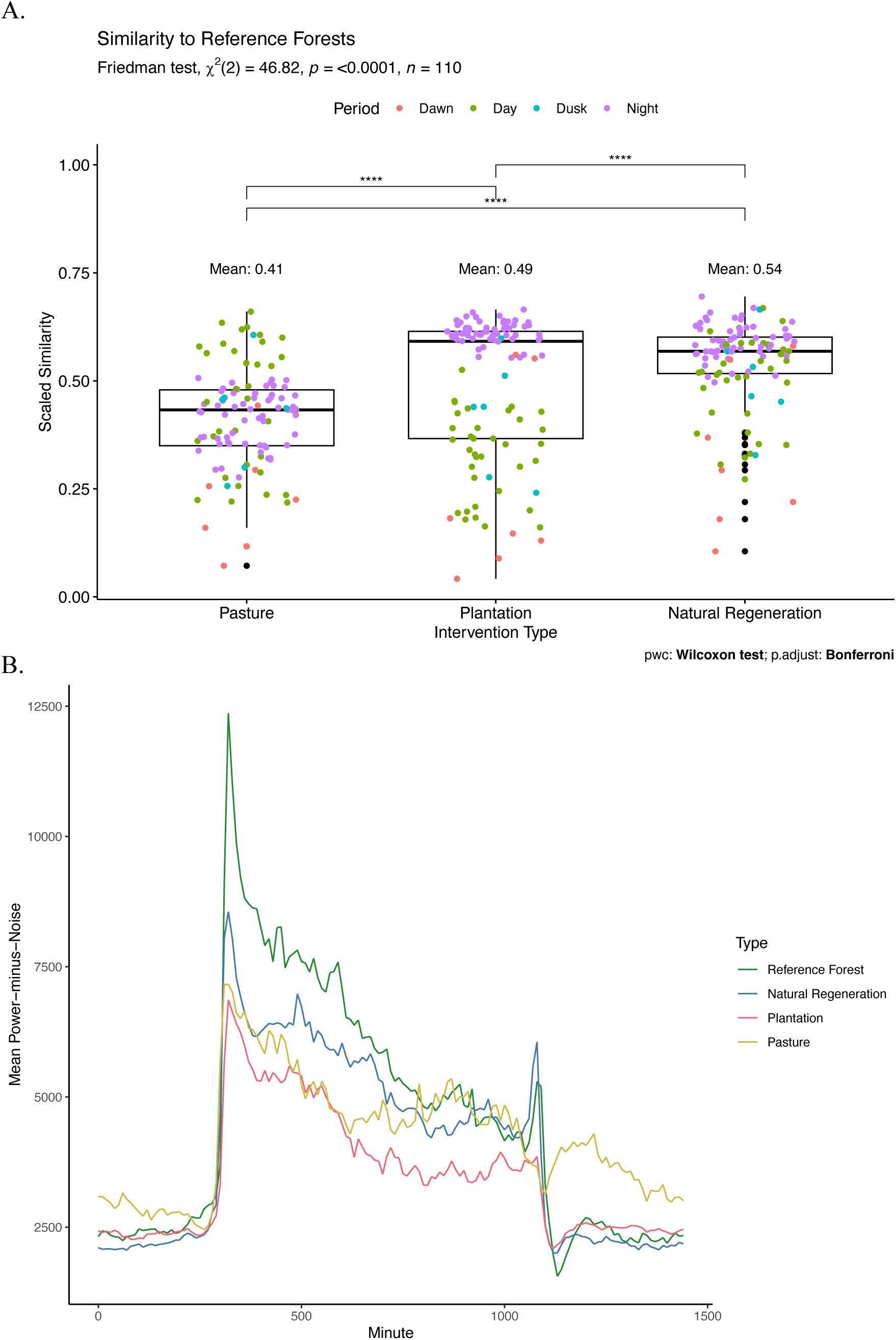
Soundscape comparisons reveal substantial recovery in PES natural regeneration sites. (A) Scaled similarity values to reference forests are shown for the remaining three land-use types, higher values indicate more shared acoustic characteristics. Box plots show the quantile ranges and median values for each land-use type. Colored points represent mean similarities for all 10-minute time bins included in the analysis, colored by the time of period of the day, while outliers are plotted again in black. Brackets indicate the results of a Wilcoxon post-hoc statistical test. (B) 24-hour cycles of SPMN for each land-use type are shown for the frequency bands in which similarities between natural regeneration sites and reference forests were highest (3 – 4 and 6 – 7 kHz).

Comparing PES sites to our baselines, we found that natural regeneration sites and reference forests shared considerable acoustic characteristics, with a mean similarity of 0.539 (Figure 3a). This was significantly larger (p < 0.01) than the similarities between either reference forests and pastures (mean similarity 0.413) or plantations (mean of 0.489). The similarities peaked during the early afternoon and particularly during the dusk chorus, where mean similarity values regularly exceeded 0.90. These similarities were strongest in frequency bands which are densely occupied by native fauna (3-4, 6-7kHz), indicating potentially shared biodiversity characteristics (Figure 3b). While natural regeneration sites also exhibited a notable similarity to pastures (mean of 0.429), these were significantly smaller (p < 0.01) than those to reference forests and could reflect the presence of generalist species common across all habitats. When comparing differences in distances to the two baselines within time-frequency intersections, we found that natural regeneration sites were, on average, 97.79% closer to reference forests than pastures. That is, for any given time and frequency, natural regeneration sites resembled reference forests almost twice as much as they did pastures.

Plantation sites within the PES exhibited intermediate similarities to both baseline ecosystems with mean similarities of 0.489 and 0.42 to reference forests and plantations, respectively. In comparison to natural regeneration sites, plantations were significantly less similar (p < 0.01) to reference forests (Figure 3a), indicating slower recovery and our data shows consistently lower levels of biological activity across the entire diel cycle. Nevertheless, plantations were significantly (p < 0.01) closer to reference forests than pastures with a mean difference in distance to the two baseline ecosystems of 79.66% towards reference forests. The similarities to reference forests were also highly variable, with high nocturnal values but similarities that dropped as low as 0.243 during the dawn chorus. Additionally, plantations did not significantly differ in their similarity to pastures compared to natural regeneration sites.

## Discussion

Despite international recognition of the need to halt and reverse the loss of biodiversity on Earth, evidence of broad restoration of terrestrial ecosystems remains scarce^44^. We used ecoacoustics to identify the ecological signals of ecosystem recovery in Costa Rica’s Payment for Ecosystem Services program across a broad geographic extent. We found that the redistributive implementation of Costa Rica’s PES program successfully moved the soundscapes of naturally regenerating forests closer to a high-integrity baseline, exhibiting mean similarities that were 97.79% closer to reference forests than to pastures (Figure 3a). These results provide tangible evidence of ecological recovery at a broad scale and provide support for equitable wealth redistribution as a financing mechanism for restoration under certain contexts.

Soundscapes contain a wealth of information that can help track changes in biodiversity and ecosystem health. Our frequency-range estimations revealed patterns that are consistent with acoustic niches^29^ occupied by distinct taxonomic classes. Specifically, mammals and amphibians largely call in the lower frequency bands, while birds and insects have highly diverse calls which extend into higher frequency ranges (Supplementary Figure 3b). However, we found humans disturb these natural acoustic dynamics directly by occupying acoustic niches (i.e., noise pollution) in lower frequencies and perhaps indirectly by disturbing ecosystems, leading to altered acoustic dynamics among land-use types. We reiterate growing calls for acoustic studies to explicitly account for multiple aspects of soundscape variation (e.g., time, frequency, amplitude) to improve ecological conclusions.

Focusing on both the time and frequency ranges indicative of natural forest communities (Figure 1), we compared soundscapes recorded from four distinct land-use types. We not only distinguish between land-uses but also investigate patterns in biological activity, which evidence suggests may correlate with patterns of ecological diversity and recovery^22,26^. In contrast to reference forests, pastures are marked by significant human influence and altered soundscape dynamics especially temporally, which indicate changes in underlying community assemblages possibly in response to the novel ecosystems these seasonally flooded grasslands provide^45^.

Across the PES sites, we found that the soundscapes of natural regeneration sites were almost twice as similar to reference forests than they were to those of pastures that represented their state prior to restoration. These results indicated substantial and widespread soundscape recovery consistent with ecological recovery, which mirrors findings for other tropical secondary forests of similar ages (30-50 years)^46,47,48^. Similarities were concentrated in the dusk chorus and in frequencies that suggest a higher rate of recovery of birds, which may be returning faster due to their dispersal abilities. Subsequent studies should use these findings as guideposts to target monitoring efforts. Plantation sites had significantly lower similarities to reference forests compared to natural regeneration sites (Figure 3a) indicating that their acoustic recovery is slower, possibly due to the lack of key components of native biodiversity^49,50^. Interestingly, plantations and natural regeneration sites did not significantly differ in their similarity to pastures, suggesting that plantations have a unique ecological trajectory that does not resemble natural forest ecosystems but is ecologically modified in ways that differ from pastures^51^. The observed results highlight the potential advantages to the PES of natural regeneration from an ecological restoration perspective^52^. Research suggests that biodiversity within plantations could be improved with different management decisions, as all the plantations in our study (and indeed most within the PES) were monocultures of non-native species planted in relatively short (10-20 year) harvesting cycles, which may contribute to the observed patterns and slowed recovery. Together, we believe these results constitute strong support for the PES as a successful intervention in recovering some aspects of ecological diversity and some of the most robust evidence to date that restoration initiatives can benefit biodiversity on large spatial scales.

Today, the PES program is widely regarded as a success and an integral part of Costa Rica’s progress in protecting its natural heritage. Having recognized the link between ecological degradation and inequality, the program is built on the need address the social challenge that underpins environmental destruction at scale. Costa Rica’s attempt to distribute wealth directly to those living in association with nature – through taxes on environmentally destructive behavior and international grants – represents a model that could be replicated and expanded globally. Weaknesses in the PES program remain and improvements are needed to achieve true equitable wealth redistribution, but studies suggest that many program participants do perceive strong socio-economic benefits^12,53^. Learning from attempts to improve Costa Rica’s PES^54^ is especially relevant given increasing environmental degradation and inequality in the 21^st^ century.

## Conclusion

We provide tangible empirical evidence for large-scale ecological recovery across PES sites in Costa Rica’s Nicoya Peninsula, following 27 years of the national Payment for Ecosystem Services program. Given that Costa Rica’s PES sites collectively cover over 200,000 hectares, we believe this is likely to be indicative of the largest recorded ecological recovery program. We show that naturally regenerated secondary forests on private lands enrolled in the PES can harbor acoustic signatures that are strikingly similar to those of protected areas, suggesting similar programs could help reverse the loss of biodiversity on a global scale. Our research emphasizes that, by prioritizing the redistribution of wealth alongside nature protection, local communities can become the stewards of biodiversity and enhance ecological health at massive scales. Our research adds to a growing body of literature that highlights the tangible links between social equity and the health of ecosystems across landscapes. Embracing these radically simple solutions to recovering biosphere integrity unlocks the possibility to simultaneously address the challenges of biodiversity loss and global inequality.

## Code availability

All the R code used for the analyses are available to reviewers from the corresponding author upon request. The code will also be made publicly accessible by the time of publication.

## Data Availability

By time of publication, all data will be available to view and download from the ETH data library platform.

## Author Contributions

GLD devised the study, carried out the fieldwork, collaborated on the data analysis and wrote the manuscript. JH heavily contributed to the data analysis, index creation and provided feedback on the manuscript. DHD and TBL helped shape the analytical approach and improved various elements of the manuscript. LKW, RC and YL provided substantial feedback on the manuscript during multiple rounds of internal revisions. CDQ, JJF, AMR, EMS, GNC, MC and ISP provided essential support in ensuring the research was relevant for local stakeholders, securing permits, extracting relevant information from government databases, and executing the fieldwork in Costa Rica. LV carried out the manual annotations. TWC guided the development of the project and collaborated closely with GLD in the writing of the manuscript.

## Methods

### Study area and site selection

The study was carried out in the Nicoya Peninsula of Costa Rica (Figure 1) between May and July of 2022. The peninsula contains both moist and tropical dry ecological zones^55^, a pronounced rainy season (May through December) and altitudes ranging from sea level to over 900m.

A total of 142 sites were initially selected from four land-use types (Supplementary Figure 1). 101 sites protected by the PES program for at least seven years were selected with support from the regional FONAFIFO office. We included sites from two PES interventions, 50 natural regeneration sites and 51 plantation sites. Natural regeneration sites are standing forests at the time of PES application. Internally the PES program refers to these as “Conservation” sites. However, no included sites were primary forests, and most have regrown on abandoned pastures^16^, making them between 30 to 60 years old. We did not encounter any PES participating landowners who had actively restored forests on their farms, therefore we refer to these sites as “natural regeneration sites”. To increase forest cover, the PES program also allows forest plantation owners to receive payments. These timber plantations are often planted with a single or few species of tree, most commonly Tectona grandis, and are harvested in 15- to 20-year cycles. As most land in the region was originally cleared for pasture, we recorded at 20 pasture sites as a benchmark to represent the state of ecosystems prior to restoration. Finally, we selected 21 primary forests or older secondary forests to represent the status of natural ecosystems outside the program. Most of these forests had been protected by the national park system (SINAC) for years to decades before the establishment of the PES program. Other sites protected by private individuals or organizations, may be in earlier stages of succession, but these number relatively few compared to the SINAC sites. Site selection, access and permission were secured in close collaboration with FONAFIFO and SINAC.

### Hardware and Acoustic Recordings

Audiomoth recorders (www.openacousticsdevices.info) versions 1.1 (n=30) and 1.2 (n=20) were used to record at the 142 selected sites. This hardware was selected for its low-cost and open-source design, which would allow other projects to replicate our methods with relatively low levels of up-front investments. We recorded continuously for six days at each site, with the aim of recording at least 120 hours per site as suggested in Bradfer-Lawrence (2019)^23^. We used a sample rate of 48 kHz and medium gain and wrote files to microSD cards in WAV format.

Recorders were placed within sites to avoid edge effects and possible interference from rivers, streams, the ocean, human settlements, or high-traffic motorways. For PES sites, each contract with FONAFIFO was considered to be an individual site and only a single recorder was placed, regardless of its size. Recorders were attached to trees at 2m height, however lack of woody vegetation in pastures required us to occasionally attach recorders to fence postings and at lower heights. We attempted a minimum distance of 700 m between recorders, to avoid recording the same acoustic event(s). Most distances exceeded 1 km (average distance between sites was 37 km), nevertheless, we retained the 7 site-pairs with geographic distances below 700m.

Canopy height was taken using a laser-range finder three times along a 10m transect centered around the recorder. PES contract ages and forest patch sizes were provided by FONAFIFO. Hardware failure or excessive acoustic interference resulted in incomplete, low-quality, or failed recordings at 22 sites, such that all further analyses include a total of 119 sites: 18 Reference Forest, 39 Natural Regeneration, 43 Plantation, and 19 Pasture sites. Recorder issues also resulted in few data points for three minutes of the day (minutes 385, 386 and 1,080), so we excluded these minutes from all analyses.

### Acoustic Processing and Index Calculation

We collected a total of 999,470 unique minutes, or nearly two years, of audio data. While many acoustic indices already exist, the complexity of these indices often make them highly context-specific and difficult to interpret, potentially leading to misunderstanding and misapplication^31^. As such, we focused on acoustic energy as this is easily comprehensible – more energy equals more sound – and readily tied to both observed soundscape patterns and underlying biological events.

Using R (version 4.1.2) and the seewave^56^ and soundecology^57^ packages we calculated a modified version of the Power Minus Noise (PMNsp) statistic as described in Towsey (2017)^35^. In brief, PMNsp is a vector derived from a spectrogram from which the modal amplitude (or energy) per frequency-band (n = 256 per recording, assuming a FFT of 512) is removed (this modal value represents the background noise or BGNsp). The de-noised spectrogram is then smoothed and the resulting maximum amplitude value per band is retained as PMNsp for every minute. While the BGNsp removal performs well in removing constant geophonies (e.g., rain and wind), it also removes some constant insect-produced noises (e.g., cicada calls). As this removal applies across the dataset and more variable insect noises are retained, we chose not to compensate for this loss of information, but we did retain BGNsp values as predictors in the models (see below). Rather than the difference between the maximum decibel value and BGNsp value for each frequency band, we took the sum of differences between amplitude values and the BGNsp across all time bins (n = 5625, each ∼0.01sec), for each band and minute to derive ΣPMN.

The interpretation of this index is straightforward, with high values indicating soundscapes with complex and variable energy levels, such that sonotypes regularly rise above the modal amplitude value. Low values indicating increasingly similar modal and max amplitude values, either because both remain low (e.g., silence) or because the soundscape is heavily completely saturated with high levels acoustic energy (e.g., rainstorms).

In most of our analyses we used 10-minute mean ΣPMN values. These 10-minute means were summed across frequencies as needed, either for a range of bands - giving a single value per time bin across all considered frequencies - or for 1 kHz at a time. Thus, our modified index gives a descriptive measure of total acoustic power per 10-minute bin and frequency range of interest.

### Index Visualization and Predictive Variable Modelling

To test the impacts of a set of predictive variables on ΣPMN we used two approaches: gradient boosted machines and linear mixed models. First, the gradient boosted machine was used to quantify the relative importance of each of 8 predictive variables: Time (10-minute bins), Annual Precipitation, Enhanced Vegetation Index, Human Modification, Elevation, Vegetation Type, Mean Canopy Height and summed BGNsp as described above for ΣPMN. We trained the model to predict ΣPMN. The index was first summed across all frequency bands and then we took 10-minute means across all sites within land-uses. The variable importance of each predictive variable was assessed by SHAP analysis^58^ (Supplementary Figure 2). To avoid overestimation of our categorical variable for land-use we included each land-use type as a binary variable (True/False) and aggregated the SHAP importance into a single variable without direction.

Second, to restrict the temporal acoustic space for our subsequent analyses, we constructed linear mixed models using per-minute ΣPMN values as the response variable. Models for each possible combination of the 7 environmental variables (same as those mentioned above apart from time) were fitted to each minute of the day and we selected the model with the lowest AIC score. Hardware identity (Gen1.1 vs Gen1.2 audiomoth recorder) was included in every model as a random factor, to account for potential technological differences. Relevant time ranges for analysis were identified by including any minute for which “land-use type” was retained in the final model. However, as most analyses are aggregated into 10-minute time bins, we included any 10-minute bin in which more than half of minute-specific models contained site-type as a predictor.

### Taxonomic Group Frequency Range Estimations

To connect observed soundscape patterns to underlying biological processes it is necessary to estimate the frequency ranges occupied by taxonomic groups of interest. While other studies have used a-priori knowledge or literature-based estimates^32^, we derived dataset-specific estimations by manually annotating 120 minutes of recordings. To avoid biases, a random number generator was used to pick minutes from sites representing the full 24-hour cycle (60 diurnal and 60 nocturnal) and land-use types (30 for each). Chosen minutes were loaded into Raven (Cornell Ornithology Lab) where every visible and audible sonotypes, or individual sound element, was carefully selected to measure the maximum and minimum frequencies. Using an independent training set of 20 minutes a lab technician (LV) was trained by authors with extensive field experience in the tropics (GLD, LW) to identify sounds as having been produced by amphibians, birds, insects, or mammals. The lab technician then annotated the 120 selected minutes. Any sounds that could not be identified with certainty by the authorship team were marked as “unknown”. We also marked and identified anthropogenic (human produced) sound and distinguished between wild and domestic mammals. The resulting frequency ranges for each taxonomic group were then exported into R where the mean frequency characteristics (mean minimum, maximum and midpoint frequency) and occupancy densities per taxonomic group were calculated (Supplementary Figure 3b). We used an ANOVA and post-hoc pairwise test to determine whether mean frequency characteristics differed significantly between taxonomic groups.

### Distributional Comparisons of Acoustic Energy

With an understanding of the frequency ranges occupied by different taxonomic groups in our dataset, we again compared the soundscapes across land-use types, now broken down into 1-kHz frequency bands. Based on our recording annotation we determined that the most biologically active frequency range in our dataset lay between 0 and 9 kHz (95% of all calls with biological origin fell within this range), which is consistent with previously published ranges and those used for index calculations (e.g., NDSI)^59^. To quantify the resulting differences between land-use types and identify the PES intervention that most closely resembled reference forest soundscapes, we employed a Wasserstein distance calculation. The Wasserstein distance, or earth-movers distance, considers two distributions and calculates the minimum ‘cost’ of moving one into the other, taken to be the amount of ‘earth’ multiplied by the distance moved. The distributions of land-use types are made up of ΣPMN summed across 1 kHz frequency bands within sites and then averaged for every minute within the 10-minute range of interest. This results in a separate distribution of points = (10 × number of sites) for each time frequency intersection. Distance metrics were calculated to every land-use type; however, we report only acoustic distances to pastures and reference forests (Figure 3a). We used a Friedman non-parametric statistical test for repeated measures to detect differences in distances across 10-minute time bins and a Wilcoxon signed-rank post-hoc test with Bonferroni correction to identify significant differences among land-use types.

These distances are reported as similarity after having applied an inverse exponential scaling of y = e^-kx^, where x is the calculated distance and k = 0.00001. Additionally, given a set of distances for any time-frequency intersection, we calculated the percentage difference in distance from PES interventions to both baseline soundscapes. We then took the mean of these percentages across all time-frequency intersections to evaluate how much closer PES sites were to one baseline than another.

## Figures

**Supplementary Figure 1:**
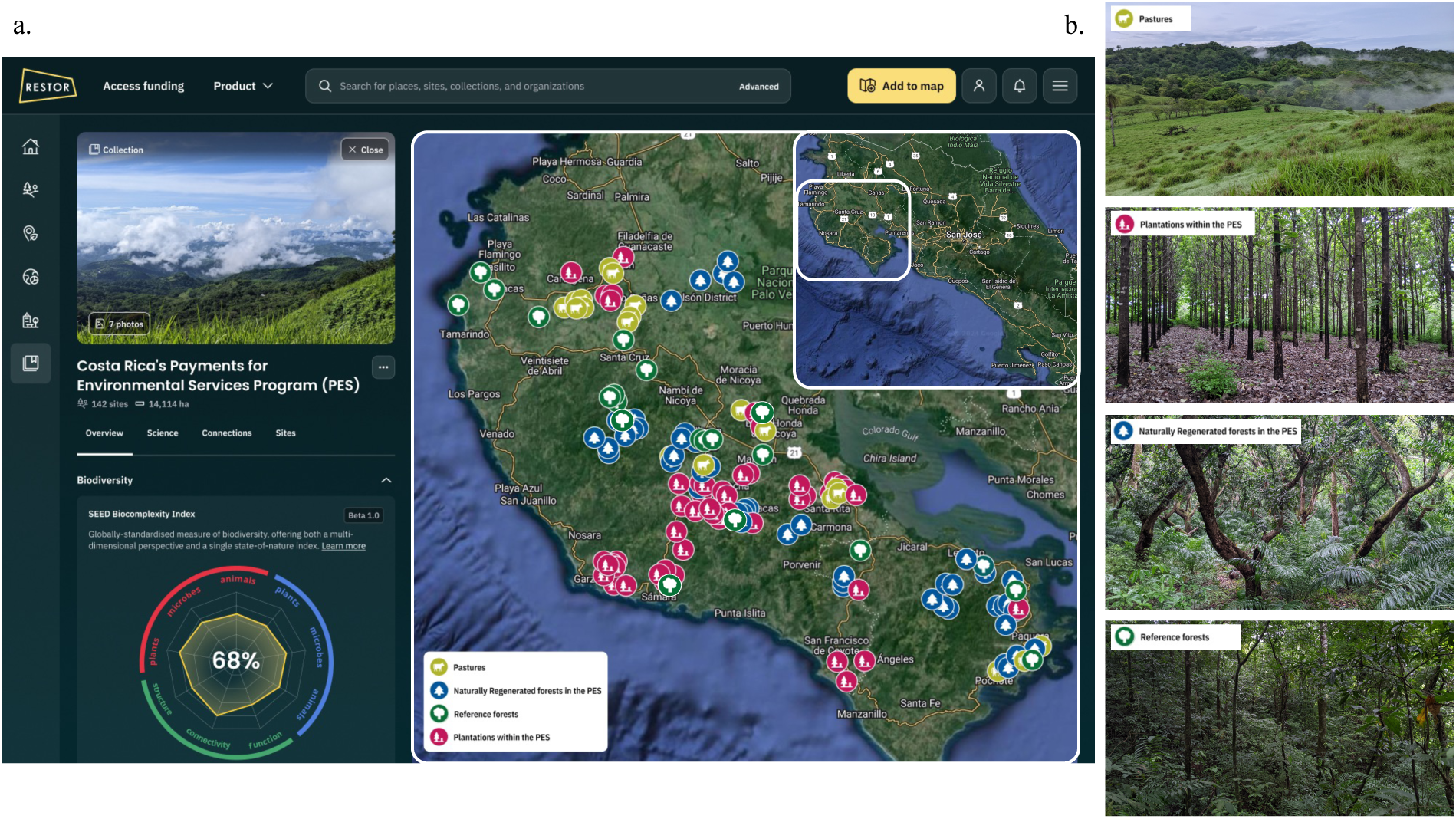
The study area in Costa Rica. (A) A map of all the recording locations in the Nicoya Peninsula as seen on RESTOR, an online open-access platform for conservation and restoration. Each point represents a single sampling location. (B) Example photos of each land-use type taken from the field during the sampling period in 2022.

**Supplementary Figure 2:**
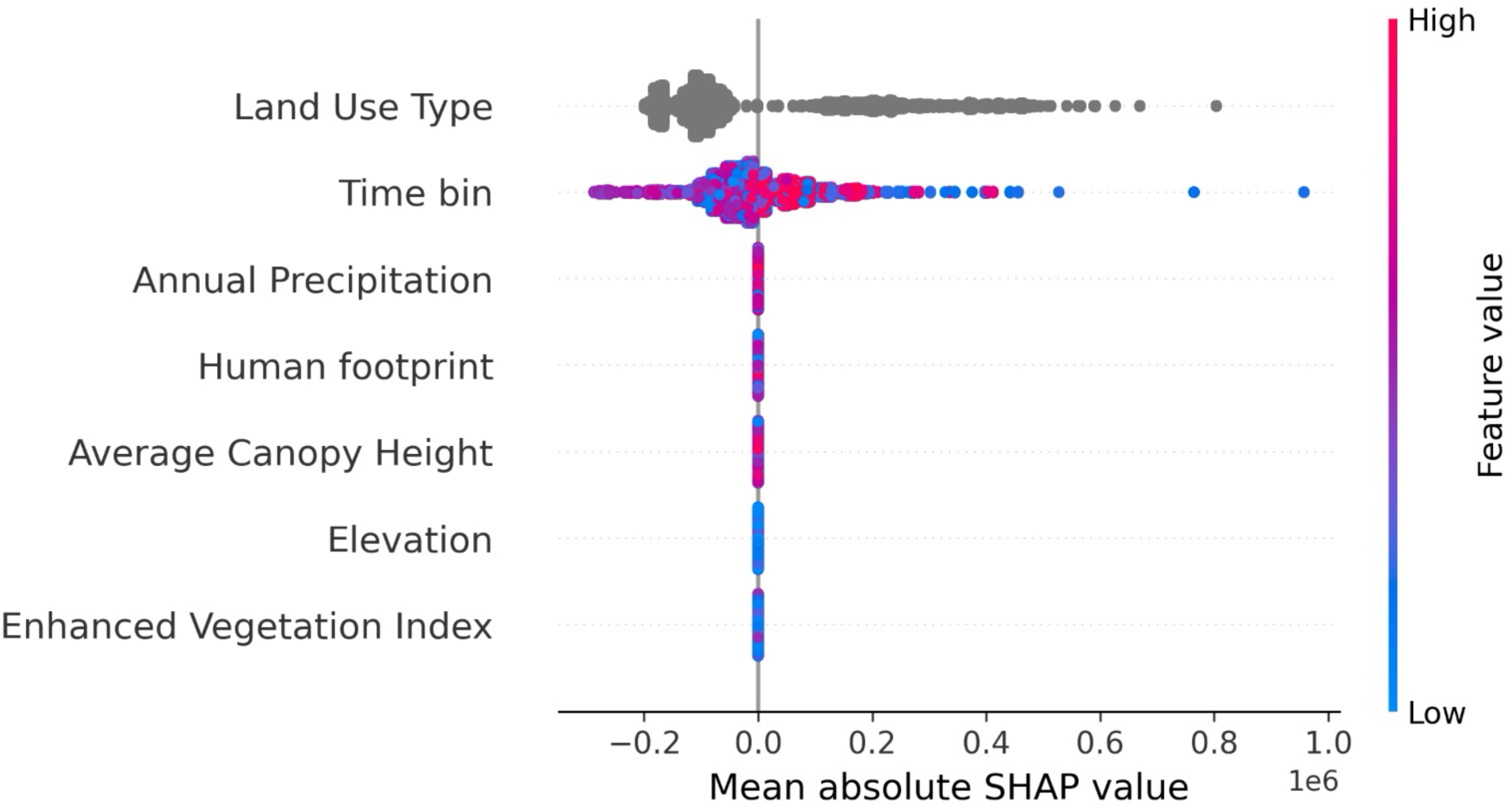
Establishing the utility of SPMN. The results of a SHAP analysis on gradient boosted machine models indicate that land-use type is the strongest predictor for variations in ΣPMN, indicating that our index captures habitat-specific acoustic dynamics.

**Supplementary Figure 3:**
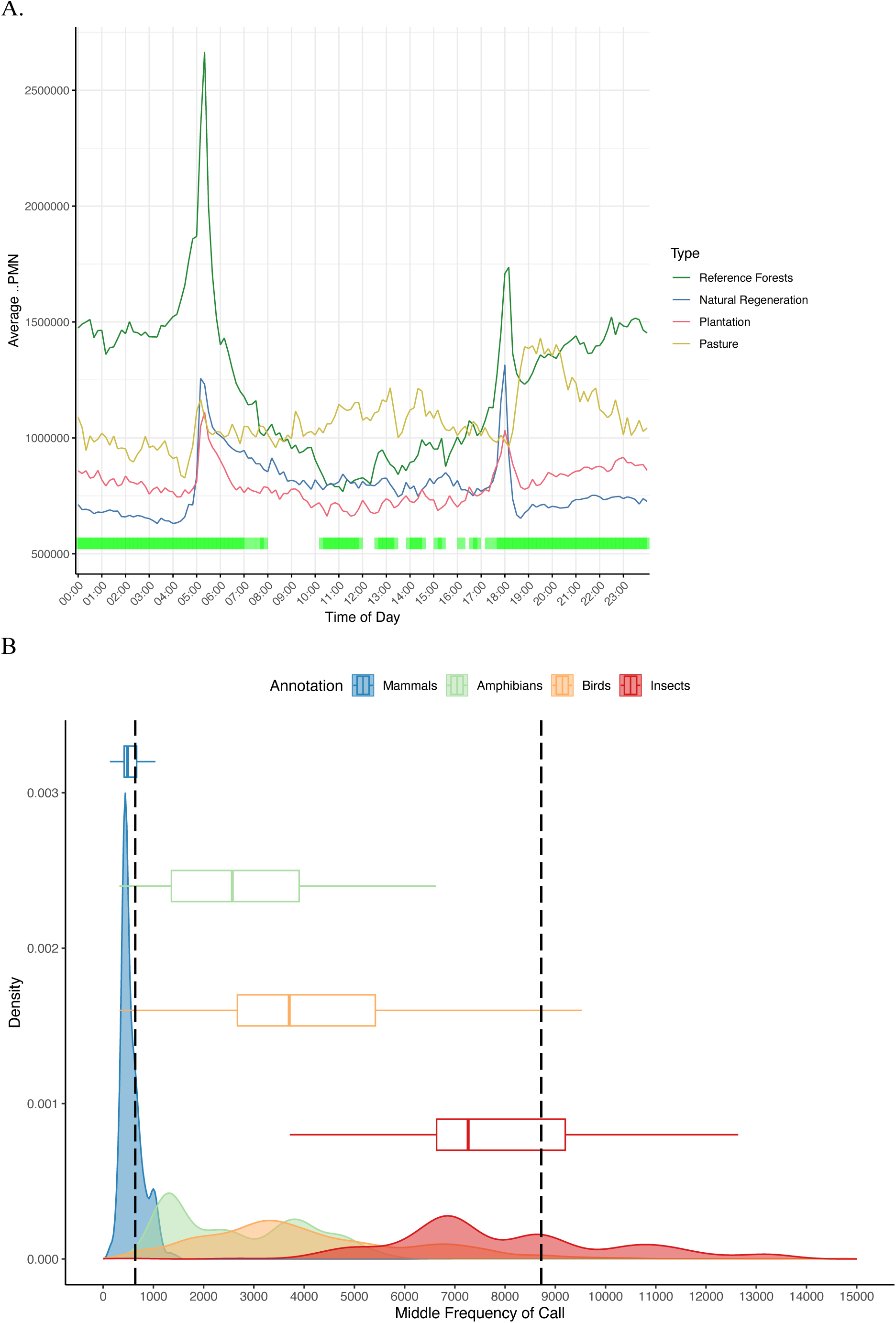
Identifying the relevant times and frequencies for analysis. Figure 2 (A) Mean 24-hour cycles of ΣPMN are plotted for each land-use type across all frequency bands. Green bars indicate the temporal ranges where land-use type is a strong predictor of soundscape differences, as identified by our linear mixed-modelling approach. (B) Density plot and boxplots for frequency ranges of four main taxonomic classes, derived from manual annotation. Black vertical lines indicate the frequencies in which 90% of biological sound occurs.

